# Metagenomic mining reveals novel viral histones in dsDNA viruses

**DOI:** 10.1101/2024.12.06.627149

**Authors:** Yang Liu, Zhuru Hou, Wanshan Hao, Shaoqing Cui, Haibo Wang, Yue Liu

**Author notes:** These authors contributed equally to this work (Yang Liu, Zhuru Hou, Wanshan Hao). Correspondence (H.W.); (Y.L.).

## Abstract

The compaction of genomic DNA into nucleosomes with the help of histones had long been considered as a fundamental feature exclusive to eukaryotic cells. However, it was recently shown that archaea and bacteria also encode histones. A complex picture has emerged with more recent discoveries of eukaryotic-like histones within one phylum of double-stranded DNA (dsDNA) viruses. Nevertheless, the extent to which other dsDNA viruses encode histones remains largely unexplored. Here we conducted a metagenomic survey of viral histones that were further clustered based on sequence and predicted structural similarities. We identified over 1,500 viral histones and histone-fold proteins, including previously undescribed proteins found in the viral class *Caudoviricetes*. Structural predictions and *in vitro* assays demonstrated that histone triplets (where three histone folds are fused) and singlets co-occurring in the same viral genome are capable of forming nucleosome-like particles. Beyond nucleosomal histone functions, our analysis revealed six types of structurally and functionally diverse viral histone-fold proteins, some of which do not have known structural or functional homologs. Altogether, our findings reveal a previously unrecognized diversity of viral histones in dsDNA viruses, expanding the known repertoire, structural diversity, and functional versatility of viral histones beyond nucleocytoplasmic large DNA viruses.

**Highlights:**

- Histone triplets and singlets co-occurring in the same dsDNA viral genomes assemble into nucleosome-like particles
- Six novel types of structurally and functionally distinct viral histone-fold proteins are identified
- The presence of viral histones is significantly positively associated with proteins containing SNF2 chromatin remodeling domain found in viral genomes

## INTRODUCTION

Histones are essential components of chromatin in eukaryotic cells that regulate DNA replication, DNA repair, and gene expression by compacting DNA and modulating the accessibility of genetic information through epigenetic modifications [1]. Over the past two decades, interest has grown in exploring the origin of eukaryotic histones and their connection to eukaryogenesis [2]. The discovery of structurally distinct histone homologs in archaea, e.g. HMfA and HMfB, has suggested that histones emerged before the divergence of archaea and eukaryotes [3–6].

Archaeal and bacterial histones differ significantly from canonical eukaryotic histones. For instance, some archaeal histones lack N-terminal tails and can form hypernucleosomes. These hypernucleosomes have the potential to form more complex structures than eukaryotic nucleosomes, which feature a histone octamer core [7]. Additionally, the archaeal histone MJ1647 from *Methanocaldococcus jannaschii* has been proposed to function as a DNA-bridging protein [8]. Similarly, the bacterial histone Bd0055 from *Bdellovibrio bacteriovorus* can form a variety of quaternary structures capable of bridging or bending DNAs [9,10]. Furthermore, a more recent systematic computational study identified over ten types of histone-fold proteins from bacterial and archaeal genomes [11], indicating a much greater diversity of histones in bacteria and archaea than previously anticipated.

Nearly all eukaryotic viruses with a double-stranded DNA (dsDNA) genome replicate their genomes in the host cell nucleus. To achieve this, some viruses (e.g., hepatitis B virus, herpesviruses, papillomaviruses, and polyomaviruses) can hijack host histones to condense the viral genome into nucleosome-containing viral chromatin, effectively mimicking the host chromatin structure and facilitating viral replication, transcription, and evasion of host immune responses [12–15]. Unlike these viruses, emerging research has revealed that a portion of large dsDNA viruses in the phylum Nucleocytoviricota, hereafter referred to as NCVs, encode histones in their own genomes [16]. For instance, computational, biochemical, and structural analyses showed that NCV histones can wrap DNAs to assemble into nucleosome-like structures, suggesting their proto-eukaryotic origins [17]. Furthermore, many NCVs also contain histone repeats where multiple histone folds are fused in a single protein (e.g., H2A-H2B-H4-H3), that can form oligomers of nucleosome-like structures resembling the aforementioned hypernucleosomes in archaea [18]. These viral histone repeats have been proposed as an intermediate evolutionary state, further suggesting that viral histones may have originated prior to the divergence of archaea and eukaryotes, potentially during eukaryogenesis [18,19]. Nevertheless, the question of whether other dsDNA viruses encode histones remains largely unexplored, impeding our understanding of histone structure, function, and evolution.

In this work, by mining of the IMG/VR metagenomic database, we identified over 1,500 viral histones, including previously undescribed histone-fold proteins in viral class *Caudoviricetes*, which primarily infect archaea and bacteria. Our structural and functional analysis revealed that *Caudoviricetes* contain predominantly bacterial-like and archaeal-like histones, as well as a minor group of eukaryotic-like histones, Moreover, *in vitro* assays demonstrated that viral eukaryotic-like histones, particularly H2B-H2A-H3 triplets and H4 singlets co-occurring in the same viral genome, can assemble into nucleosome-like particles. Furthermore, in addition to nucleosomal histones, we discovered six novel types of viral histone-fold proteins. Whereas some types exhibit structural similarities to known bacterial or archaeal histones, many do not have known structural or functional homologs. Finally, we showed that viral histones frequently positively associate with specific putative chromatin-related proteins (CRPs) in *Megaviricetes*, including those containing the SNF2 chromatin remodeling. These findings highlight a previously unrecognized diversity of viral histones, of which future structure-function studies will provide new insights into understanding virus-host interactions.

## RESULTS

### Identification of viral histones in double-stranded DNA Viruses

In order to identify viral histone proteins from metagenomic assemblies, we employed Hidden Markov Model (HMM) profiles of eukaryotic (H2A, H2B, H3, and H4), bacterial (*Bdellovibrio bacteriovorus* Bd0055), and archaeal (*Methanothermus fervidis* HMfA/B) histones derived from previous studies [9,18]. A profile-specific HMM search was conducted against 5,576,197 high-confidence viral genomes from the IMG/VR database [20]. This analysis yielded 3,301 raw hits, of which 966 proteins, with domain similarity e-values below 1×10^-5^. These histone proteins were identified primarily in the viral classes *Megaviricetes* and *Caudoviricetes*, and only marginally in *Naldaviricetes* and *Pokkesviricetes* (Figures 1A and S1A). Whereas members of *Megaviricetes* belong to Nucleocytoviricota viruses that typically infect eukaryotic hosts like amoebae, algae, insects, and animals [21], viruses within *Caudoviricetes* are tailed bacteriophages that mainly infect bacteria and archaea [22]. We next clustered the identified viral histones based on both structural and sequence similarities (see method), and assigned cluster annotations (eukaryotic-, archaeal-, or bacterial-like) according to the respective histone HMM profiles. The viral histones most closely related to eukaryotic reference histones H2A, H2B, H3 and H4 were identified primarily in *Megaviricetes* (Figures 1B-C). In contrast, the most abundant histones found in *Caudoviricetes* primarily had the closest match to archaeal and bacterial reference histones (Figures 1B-C). In particular, the 84 bacterial-like histones identified in our analysis have not previously been reported to be virally encoded. Furthermore, viral host predictions revealed that members of *Caudoviricetes* containing bacterial-like and archaeal-like histones were predicted to infect the phyla Pseudomonadota (bacteria) and Methanobacteriota (archaea), respectively, consistent with the source phyla of the reference histones used in our HMM-based search (Figure S2). This result suggests that *Caudoviricetes* may have acquired histone genes from their hosts. Overall, whereas all bacterial-like histones are quite different from eukaryotic-like and archaeal-like histones, archaeal-like histones show a close similarity to eukaryotic-like histones, supporting the proposed archaeal origin of eukaryotic histones (Figure 1C) [3,19].

**Figure 1.**
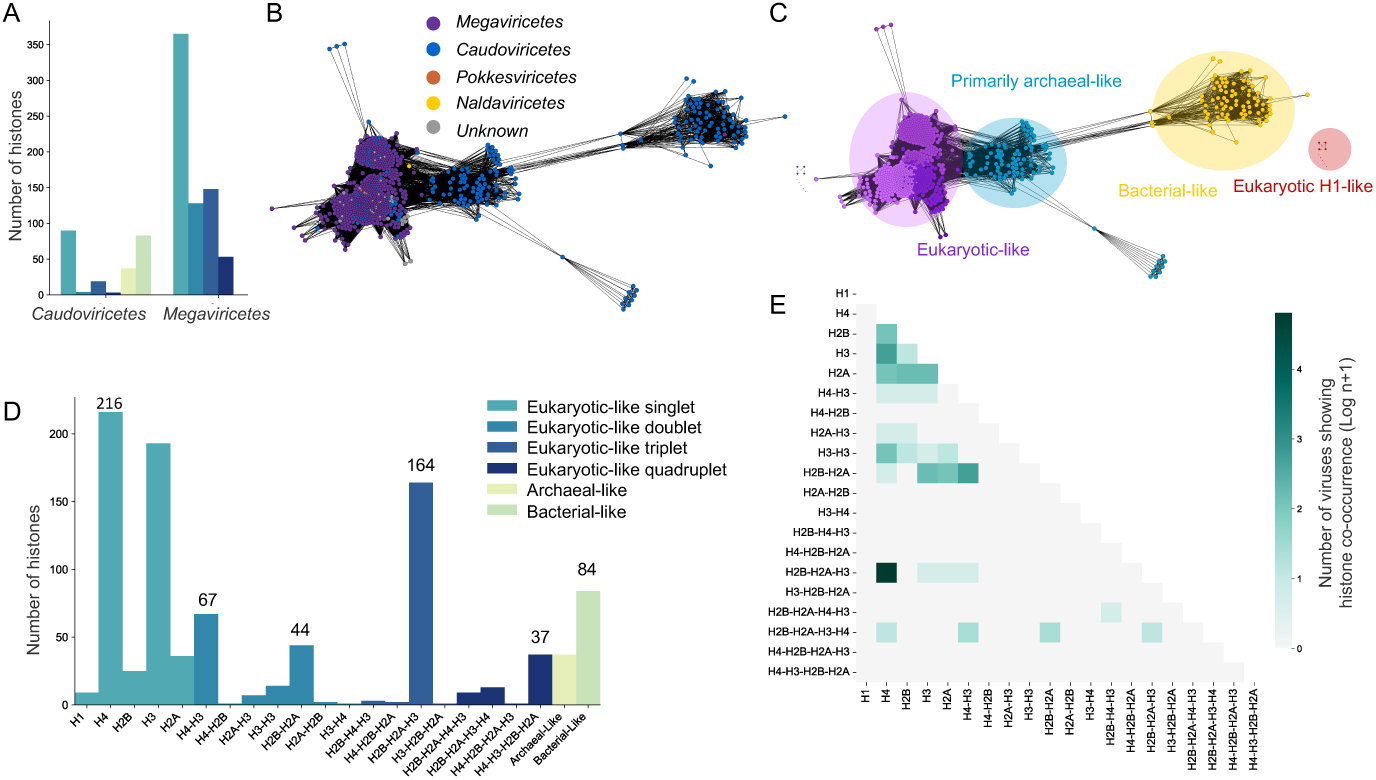
An HMM-profile based metagenomic survey of the IMG/VR database identifies diverse viral histones. (A) Abundance of different viral histone types across two viral classes, where histone types are annotated by their corresponding HMM profiles and colored as in (D). (B) Viral histone similarity network colored based on viral classes. Each node indicates a histone, and the edge represents the similarity between two viral histones. (C) Viral histone similarity network (identical to that in (B)) colored according to sequence and structural similarities. The eukaryotic-like histone cluster contains three well-connected sub-clusters. (D) Histone fold type and their abundances. (E) The co-occurring rates of eukaryotic-like histone pairs in same viral genomes.

A more detailed analysis showed that the distribution of histone protein lengths observed in *Caudoviricetes* is comparable to that in *Megaviricetes*, showing two main peaks (∼165 and ∼380 amino acids) in abundance (Figure S1B). Many histones within the second peak differ from the canonical eukaryotic histone singlets (i.e., H2B, H2A, H3, and H4). Instead, these eukaryotic-like histones can be annotated as histone repeats, including doubles, triplets, and quadruplets, based upon their corresponding position-specific HMM profiles (see Methods). We found that singlets H4 and H3, doublet H4-H3 and H2B-H2A, triplet H2B-H2A-H3, and quadruplet H4-H3-H2B-H2A are among the most abundant types of histones (Figure 1D). Viral genomes that encode histone triplets and quadruplets showed a significantly small number of total genes compared to those containing histone singlets or doublets (p<0.05, Mann-Whitney U test) (Figure S1C). In addition, viral genomes containing histone triplets and quadruplets showed more histone genes per genome (Figure S1D).

Consistent with previous studies showing that multiple pairs of histone types frequently co-occur in the same viral genomes [23,24], we also found similar patterns of co-occurring histone pairs. Among these, the co-occurrence of viral H2B-H2A with H4-H3 and that of viral H2B-H2A-H3 with H4 have the highest counts (Figure 1E). It was previously shown that H2B-H2A-H3-H4 quadruplets as well as co-occurring H2B-H2A and H4-H3 doublets can similarly wrap DNAs or form nucleosome-like particles, indicating an evolutionarily conserved functional requirement for the presence of all four histone singlets in nucleosome formation as in eukaryotes. Thus, we hypothesize that an H2B-H2A-H3 triplet and an H4 singlet found in the same viral genome could similarly form nucleosome-like structures.

### Co-occurring viral H2A-H2B-H3 triplets and H4 singlets are capable of nucleosome formation

To test this hypothesis, we used AlphaFold3 to predict the ability of nucleosome formation for various combinations of viral histones in the presence of a model double-stranded Widom 601 DNA (W601) (Figure 2A). Predicted structures of H2A-H2B-H3 triplets and H4 singlets co-existing in the same genome, when complexed with the model DNA, closely resemble known eukaryotic and viral eukaryotic-like nucleosomes. These structures featured a DNA-wrapping core composed of eight histone-fold domains, which consists of two copies of H2A-H2B-H3 triplet and H4 singlet. Notably, some histone triplets show potentially disordered loops that could provide sites for epigenetic modifications and highly acidic, long C-terminal helices that may have a role in epigenetic regulation (Figure 2A).

**Figure 2.**
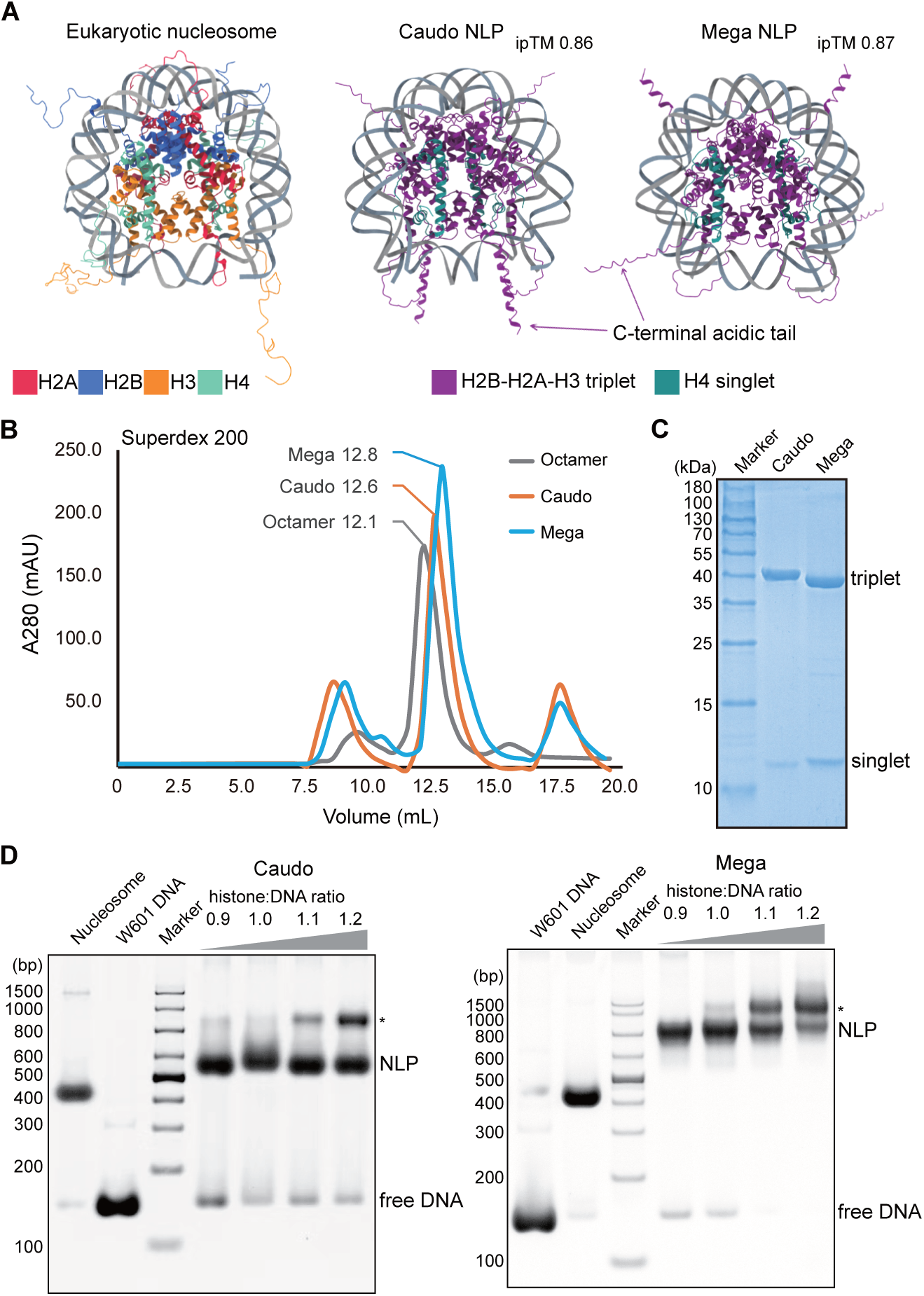
Co-occurring pairs of viral histones assemble into nucleosome-like particles (NLPs). (A) AlphaFold3 predicted structures of viral NLPs resemble that of the canonical eukaryotic nucleosome core particle (PDB ID: 1KX5). ipTM: interface predicted template modeling score. (B) Size exclusion chromatography (Superdex 200 10/300) profiles of co-expressed histone triplets and singlets from a representative virus of *Caudoviricetes* (Caudo) and *Megaviricetes* (Mega) and of the eukaryotic histone octamer, respectively. Retention volumes of the major peaks are denoted. (C) Coomassie blue-stained SDS–PAGE gel of peak fractions from co-expressed viral histones purification. Molecular weights of marker are labeled on the left. (D) Electrophoretic mobility shift assays for histones from representative *Caudoviricetes* (Caudo) and *Megaviricetes* (Mega) in combination with 145 bp Widom 601 DNA. Histone to DNA molar ratios are noted. Free Widom 601 DNA and assembled eukaryotic nucleosome on the left are used as control. Positions of formed NLPs are denoted. Higher molecular-weight assemblies, as indicated by an asterisk, are present in conditions with high histone-to-DNA ratios.

We next co-expressed and purified the H2B-H2A-H3 triplet and H4 singlet from a representative virus of *Caudoviricetes and Megaviricetes* in *E. coli*. In both cases, the size exclusion chromatography profile of co-expressed histone proteins indicates that histone triplets and singlets from the same viral genome form a complex like eukaryotic histone octamer (Figure 2B). The retention volume of around 12-13 ml on Superdex 200 10/300 column (Cytiva) corresponds to a molecular weight of around 200 kilodalton (kDa), implying that two copies of each histone triplet and singlet exist in the quaternary structure (Figures 2B and 2C). When combined with the 145 bp W601 nucleosome positioning sequence DNA and employing the classic salt-gradient dialysis nucleosome reconstitution protocol [25], H2B-H2A-H3 triplet and H4 singlet form defined nucleosome-like particles (NLPs), as evidenced by electrophoretic mobility shift assays (Figure 2D). However, viral NLPs migrate slightly slower than the canonical eukaryotic nucleosome core particle (NCP) in the gel, indicating that they might be less compact than eukaryotic NCP (Figure 2D). The presence of multiple slowly migrating species in viral NLPs suggest that they could potentially form higher order assemblies (Figure 2D).

### Identification of structural and functionally diverse viral histone-fold proteins

Beyond the role in nucleosome formation, histones have been proposed to play a wide range of other functional roles across domains of life [18]. We performed functional domain annotation for the aforementioned 966 viral histones based on their predicted 3D structural information (see Method). When compared with eukaryotic-like histones, bacterial-like histones showed much less prominent structural homology to canonical eukaryotic histones (Figure S3), suggesting that they may represent histone-fold proteins that serve distinct functions. To explore the diversity of viral histones further, we performed a HMM-based metagenomic survey by using profiles of histone-fold proteins in bacteria and archaea as previously defined [11]. A total of 289 viral no-redundant (out of 539) histone-fold proteins were identified from the *Caudoviricetes* class, and they were grouped into seven clusters based on their normalized embedding– and sequence-based similarities (Figure 3A). Consistent with the clustering results, structural comparisons showed that proteins within the same cluster exhibited higher structural similarities than proteins found between two clusters did. (Figure 3B). We describe the seven clusters of viral histone-fold proteins in details below.

**Figure 3.**
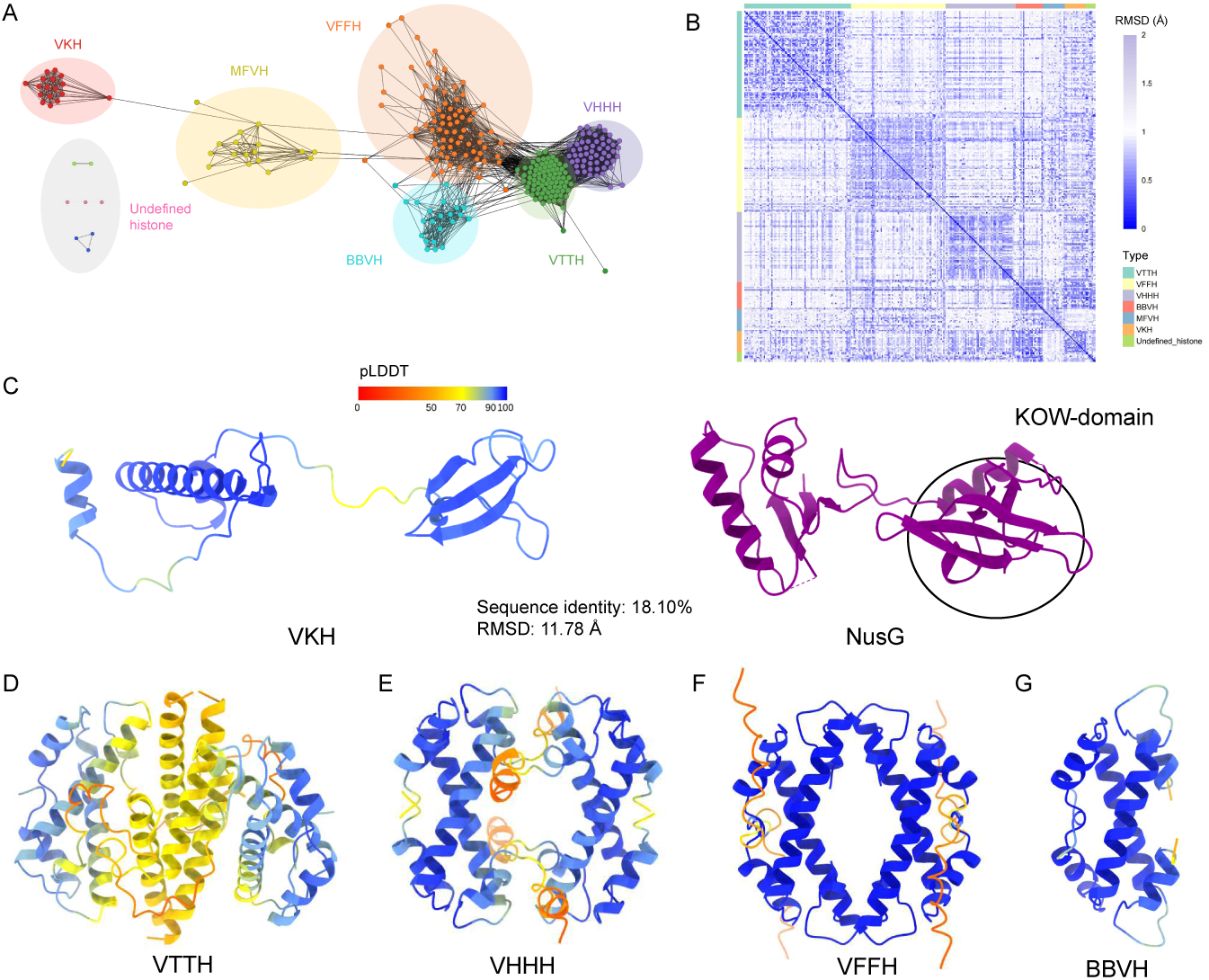
Six types of structurally and functionally diverse histone-fold proteins identified in viral genomes. (A) The structure– and sequence-based similarity network of viral histone-fold proteins. Six novel clusters are named as follows: VTTH, viral tail-to-tail histone; VFFH, viral face-to-face histone; VHHH, viral head-to-head histone; BBVH, *Bdellovibrio bacteriovorus* Bd0055-like viral histone; MFVH, *Methanothermus fervidus* Hmf-like viral histone; VKH, viral KOW histone. (B) Inter-cluster and intra-cluster structural comparisons between different histone-fold proteins. Darker colors indicate a greater structural similarity between the two proteins. (C) The AlphaFold2 predicted monomeric structure of a representative VKH in comparison with the crystal structure of NusG (PDB ID: 5TBZ). (D-F) The tetrameric structures of a typical VTTH (D), VHHH (E) and VFFH (F) as predicted by AlphaFold2. (G) The dimeric structure of a typical BBVH as predicted by AlphaFold2. These structures are colored by their pLDDT value for each residue.

Viral KOW histones (VKHs) form a previously undescribed cluster of histone-fold proteins. VKHs contain a histone fold connected by a long loop to a 27-amino-acid Kyrpides-Ouzounis-Woese (KOW) domain (Figure 3C). The KOW domain is a known RNA-binding motif, found in a variety of ribosomal proteins, the essential bacterial transcriptional elongation factor NusG, and eukaryotic transcriptional elongation factor Spt5 [26]. This domain promotes interactions with nucleic acids, ribosomes, and transcription regulatory factors such as termination factors by forming a β-barrel fold, and may be crucial for transcriptional regulation [27,28]. We found that the KOW domains in VKHs are structural homologs of NusG (RMSD: 3.60 Å) rather than sequence homologs (about 18% amino acid sequence identity with NusG). Furthermore, to our best knowledge, the KOW domain has not previously been identified to be associated with any histones or histone-fold proteins. It is probable that VKHs may be capable of binding both DNAs and RNAs via a combination of histone and KOW folds, likely playing a role in coupling multiple processes in gene regulation.

Viral tail-to-tail histones (VTTHs) represent the most abundant cluster, featuring an N-terminal α3 histone fold, named after a truncated α3 helix, that is linked to a long C-terminal helix by a flexible loop (Figure 3D). This α3 histone fold includes three conserved lysine residues (K6, K10, and K52), which may contribute to DNA binding. The C-terminal α-helix might be responsible for the formation of a stable VTTH homotetramer that could similarly function to bridge DNAs as does a previously identified histone, A0A2E7QIQ9 [11]. However, in contrast to A0A2E7QIQ9 that shows a more extended conformation as a tetramer, the connecting loop in VTTHs is generally shorter, resulting in a much more compact tetramer. This conformational difference suggests that VTTHs may adopt flexible and variable conformations that could facilitate DNA bridging by modulating the distance of the connecting DNAs (Figure S4).

Viral head-to-head histones (VHHHs) form another cluster of histone-fold protein with an N-terminal α3 histone fold flanked by a flexible loop (seven amino acids long). This α3 histone fold retains some conserved residues (K17, K18, K58, and R59) that could be important for binding DNAs. However, unlike VTTHs and A0A2E7QIQ9 [11], VHHHs lack the long C-terminal α-helix required for tetramer formation such that these proteins cannot form a stable DNA-bridging tetramer (Figures 3E and S5). Instead, they show a head-to-head tetrameric structure that may confer the ability of bending DNAs, with the help of their conserved N-terminal DNA-binding sites.

Viral face-to-face histones (VFFHs) closely resemble previously reported face-to-face histones in bacteria and archaea [11], and are similarly capable of forming stable tetramer structures (Figure 3F). However, VFFHs typically possess an N-terminal tail ranging from 5 to 40 amino acids in length, which are absent in previously identified face-to-face histones. The N-terminal tail contains many positively charged lysine residues, which may be involved in binding DNA and facilitate the DNA-bending function of VFFHs (Figure S6).

*Bdellovibrio bacteriovorus* Bd0055 like viral histones (BBVHs) share a high structural similarity with the bacterial histone Bd0055 [10]. They are predicted to form stable dimer structures like Bd0055 rather than a tetramer as in VFFHs. BBVHs differs only slightly from Bd0055 in that they feature a helix rather than a flexible loop at the C-terminus. Given their structural similarities, BBVHs could potentially condense DNAs using a similar DNA-bending mechanism as Bd0055 does [10] (Figure S7). In addition, there are some undefined histone-fold proteins that form a separate cluster, including several DNA-bridging histones, characterized primarily by their RdgC and coiled-coil structural domains [11] (Figure S8).

### The association between viral chromatin-related proteins and histones

In addition to canonical histones, a plethora of chromatin-related proteins (CRPs) play key roles in chromatin biology that are indispensable for epigenetic regulation of cellular homeostasis. We employed HMM profiles of functional domains derived from known CRPs to search for putative CRPs in high-confidence viral genomes in the IMG/VR database [20]. These chromatin-related domains included chromatin readers, chromatin remodelers, histone chaperones, domains responsible for histone (de)acetylation and (de)methylation, including polycomb repressive complexes (PRCs). Our results show that proteins with these domains were largely identified in the viral classes where histones were discovered, such as *Megaviricetes*, *Caudoviricetes*, and *Pokkesviricetes* (Figure S9). When compared with viral genomes where no histones were identified, histone-encoding viral genomes showed higher mean counts of putative CRPs (Figure 4A). Specifically, proteins with the SNF2 chromatin remodeling domain were the most abundant hits in histone-containing viral genomes (Figures 4A-B, S9 and S10), followed by those with domains involved in histone methylation and acetylation (Figures 4A, S9 and S10). In particular, proteins with domains involved in histone methylation often co-occur with H1, H3, H2B-H2A, and H2B-H2A-H4-H3 histones but not with archaeal-like, VTTH, VFFH, VHHH histones(–fold) proteins (Figure 4B).

**Figure 4.**
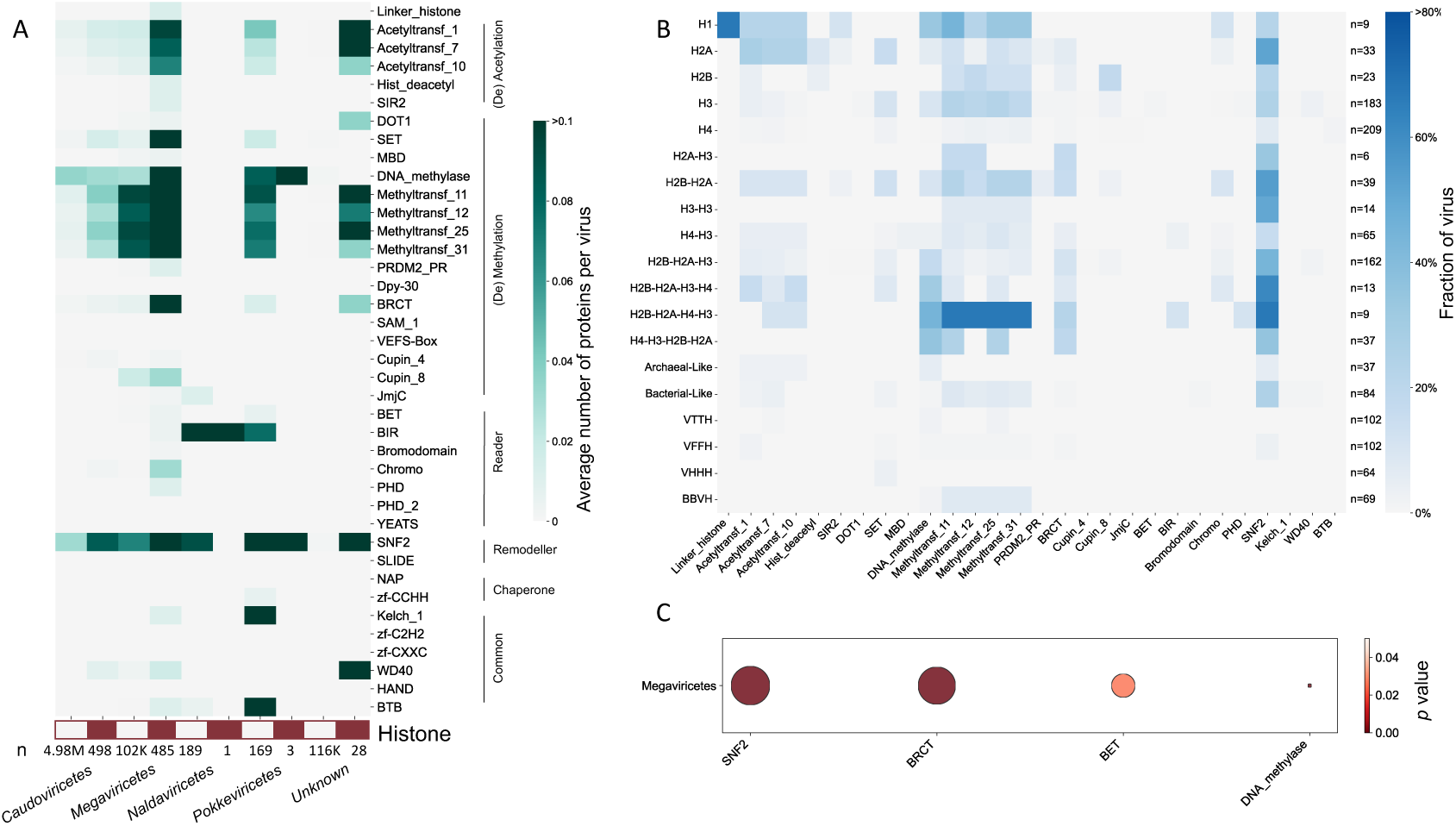
The association between histone presence and abundance of chromatin-related domain-containing proteins in viral genomes. (A) The average number of putative chromatin-related proteins (CRPs) in viral genomes for different viral taxonomic classes. The bottom red boxes indicate viral genomes with at least one viral histone, whereas the white boxes indicate genomes where no histones are identified. The letter “n” indicates the total number of viruses included in corresponding columns. All chromatin-related domains are classified into (de)acetylation, (de)methylation, reader, remodeler, chaperone, and common categories. (B) The proportion of histone-containing viral genomes that have at least one corresponding chromatin-related protein among all corresponding putative histone containing viral genomes. (C) The statistically positive correlation between the number of putative CRPs and the presence of histones in viral genomes. A positive significant correlation is colored red (*p* < 0.05). Circle sizes indicate the number of significant observations from 100 independent random samplings per domain.

We next employed a generalized linear model (GLM) to test the association between the presence of viral histone(s) and the abundance of CRPs in the same viral genomes (see Method). Overall, *Megaviricetes*, but not *Caudoviricetes*, showed a class-specific statistically significant positive correlation between the presence of histones and the number of chromatin-associated proteins (GLM test, Benjamini-Hochberg *p* adjustment, mean effect size of 0.45, and minimum *p* values of 3.95e^-10^). A detailed examination of each CRP category revealed that in *Megaviricetes*, the presence of histones was significantly positively correlated with the SNF2 chromatin remodeling domain, BET chromatin reader domain, BRCT domain, and DNA methyltransferase (DNA_methylase) domain (minimum *p* < 0.05, Figures 4C and S11).

## DISCUSSION

Our metagenomic mining identified 1,505 viral histones and histone-fold proteins from the IMG/VR database. In particular, those found in *Caudoviricetes* have not previously been described. Notably, we discovered a collection of archaeal-like, bacterial-like and even eukaryotic-like histones in *Caudoviricetes*, a class of tailed dsDNA bacteriophages primarily infecting archaea and bacteria. While our analysis highlights a structurally and functionally diverse set of bacterial-like viral histones, the eukaryotic-like histones identified are structurally and functionally similar to the canonical nucleosomal histones, as evidenced by our observations that viral histone triplets and their co-occurring singlets assemble into nucleosome-like particles. Additionally, we observed a positive correlation between the presence of histones and the abundance of proteins containing the SNF2 chromatin remodeling and BET chromatin reader domains in viral genomes within *Megaviricetes*. These findings highlight a previously unrecognized diversity of virus-encoded histones and would be instrumental for structural and functional characterizations of viral histones.

Consistent with annotations according to HMM-profiles, a majority of histones found in *Caudoviricetes* were bacterial-like or archaeal-like histones based upon structural homology with known bacterial or archaeal histones [9,29]. Sequence– and structure-based clustering enabled us to explore the structural and functional characteristics of these viral histone-fold proteins in greater details. Among these, some clusters (e.g., BBVHs, VFFHs, and *Methanothermus fervidus* Hmf like viral histone) share highly similar predicted monomeric structures and/or function-associated oligomerization forms with bacterial or archaeal histone-fold proteins lacking experimentally confirmed functions, most of which were recently predicted to bend or bridge DNAs instead of wrapping DNAs in a nucleosome-like fashion [11]. This suggests that *Caudoviricetes* might have gained these viral histones from their hosts during infection on the evolutionary timescale. However, other clusters of viral histones (e.g., VHHHs, VTTHs, and VKHs) show prominent differences in both predicted structure and putative function from their respective reference histone HMM profiles, according to which these viral histones were identified. Thus, the deletion or addition of structural elements, such as tails, helices or domains unrelated to the histone fold, during evolution could give rise to a diversity of novel histone-fold proteins, contributing to the complexity of histone structure, oligomerization, and function.

*Caudoviricetes* also encode many eukaryotic-like histones, including multiple types of histone repeats. In particular, a few eukaryotic-like histone singlets and triplets in *Caudoviricetes* share high sequence and structural similarities with their respective counterparts found in *Megaviricetes* (Figure 1C). A structure-based phylogenetic tree of histone triplets (19 in *Caudoviricetes* and 148 in *Megaviricetes*) revealed that histone triplets in *Caudoviricetes* do not form an independent lineage. Instead, they were often found adjacent to various *Megaviricetes* histone triplets (Figure S12). While we cannot completely rule out the possibilities that some *Megaviricetes* may have been mis-assigned as that of *Caudoviricetes* in the IMG/VR database due to inherent limitations of metagenomic assembly or their close evolutionary relationships [30], two possible explanations could account for the identification of eukaryotic-like histones in *Caudoviricetes*. One the one hand, the *Caudoviricetes* may have obtained eukaryotic-like histones from *Megaviricetes* through their host-host interactions, as both *Caudoviricetes* and *Megaviricetes* share similar habitats (e.g., seawater). One the other hand, some members of *Caudoviricetes* may have eukaryotic hosts, and their ancestors could have acquired histones from these hosts during the course of evolution.

A recent report by Irwin et al. proposed the hypothesis that histone triplets and quadruplets are subject to reversible genetic composition at the evolutionary level [18]. The triplets may serve as an intermediate state in the formation of histones or as a substrate for hydrolase cleavage. Given that nucleosome formation necessitates all four types of single histone folds, triplets may represent a relatively stable storage state for viral histones, offering a probable explanation for the observed high abundance of histone triplets in viral genomes (Figure 1D). Furthermore, H4 is one of the most evolutionarily conserved histones, exhibiting minimal functional diversity among all histone-fold containing proteins [31]. While this high level of conservation ensures the functional stability of H4 histone, it also limits its capacity to form novel combinations with other histones. In contrast, the variability and flexibility of H2A, H2B, and H3 may enable them to form diverse combinations more readily.

Previous studies have not observed a significant association between the presence of histones and the abundance of putative CRPs in viral genomes [1,18]. In contrast, our GLM analysis showed a viral class-specific statistically significant positive correlation in viral genomes of *Megaviricetes* when considering all categories of chromatin related domains. Specifically, several individual categories of domains were significant enriched, including SNF2 chromatin remodeling domain, BET chromatin reader domain, BRCT domain, and DNA methyltransferase (DNA_methylase) domain. It is conceivable that virally-encoded putative CRPs may have co-evolved with viral histones to affect the course of viral infections within a host cell. On the one hand, viral putative CRPs could specifically regulate viral histone or chromosome functions such as viral genome packaging and/or viral gene expression, likely via their putative chromatin remodeling, histone modification, or reader activities. On the other hand, viral CRPs could modulate virus-host interactions via alterations of the host nucleoid or chromatin architecture and functions. Viral CRPs with an SNF2 domain may facilitate changes in nucleosome positioning on the viral and/or host genome, tuning expression levels of related genes during viral infections.

In the present work, novel viral histones and histone-fold proteins were identified and characterized based on protein sequence and structural similarities. This illustrates the power of deep learning-based computational tools, such as FoldSeek and protein language models, that integrate protein structure information in homology detection and functional annotation. Nevertheless, the true repertoire and functional diversity of viral histones, including many functionally unknown variants, remains to be determined. We expect that future research on the biology of viral histones, fueled by the unprecedented availability of (meta)genomic data, state-of-the-art computational tools, and advanced approaches for structure-function analyses, will further illuminate the structure, function, and evolution of histones and their associated proteins.

## METHODS

### Viral histone identification

We used the Integrated microbial Genomes & Viral Research 4 database (IMG/VR 4) release 2022-12-19_7.1 [20] to identify all potential viral histones. The IMG/VR database comprises viral genomic assemblies identified using Genomad [32], with the majority of viral genomes originating from the Integrated Microbial Genomes & Microbiomes (IMG/M) database [33]. The high-confidence viral genome protein predictions together with their corresponding viral metadata were downloaded from FTP site. Hidden Markov Models (HMM) profiles of eukaryotic H2A, H2B, H3, H4 and archaeal histones were obtained from a recent report by Irwin et al. [18]. Bacterial histone sequences from a previous publication [9] were used to build a bacterial histone HMM profile using mafft-linsi (L-INS-I algorithm, default options, MAFFT v7.475) [34] and hmmbuild 3.1b2 (http://hmmer.org) using sequence alignment.

Each of the histone HMM profile was used to search IMG/VR high confidence proteins using HMMER 3.1b2 (http://hmmer.org). The potential histone sequences with domain e-values greater than 0.00001 were filtered out. The taxonomy information was mapped using the IMG/VR metadata. The searched histone types were assigned with their corresponding HMM profile. To determine the identity of each histone repeat, the H2A, H2B, H3, and H4 domain hits from the HMM results were initially sorted by their coordinates in ascending order. The algorithm then iteratively compared the coordinates of each domain hit with the following one. If any two HMM profile hits overlapped by more than 30% of their coordinates, the hit with the higher e-value (less significant) was removed, and its coordinates were excluded from further comparisons. In the cases where a protein did not match any of the H2A, H2B, H3, or H4 domains, it was assigned to the corresponding archaeal or bacterial histone profiles it matched. This process was applied independently to each identified histone. The Mann-Whitney U test was used to determine differences in total number of viral proteins between different types of histone-containing viruses.

To identify histone-fold proteins in viruses, we employed prokaryotic histone-like proteins HMM profiles procured by Schwab et al. [11], including bacterial_dimer, ZZ, phage_histone, coiled-coil, RdgC, IHF, face_to_face. Next, the HMM search was conducted against the IMG/VR 4 high-confidence viral database and the PhageScope database (version 1.3) [35]. Only those HMMER search results with an e-value below 0.001 were retained for further analysis.

### Prediction of histone and nucleosome structures

For 3D structural predictions of viral histones, ColabFold v1.5.2 [36] and AlphaFold v2.3.5 [37] were used, with parameters (use_templates=False, custom_template=‘none’, use_amber=False, num_recycles=‘auto’, recycle_stop=‘auto’, msa_mode=‘mmseqs2_uniref_env’, custom_msa=‘none’). Multimer v3 was utilized in the cases of homo-oligomer structure prediction. For each protein, the highest ranked atomic model out of five predicted by Alphafold2 was retained for further analysis.

To predict potential nucleosome formation for each viral candidate (*Megaviricetes* IMGVR_UViG_3300010430_005728 histone ID: Ga0118733_100000053190 and Ga0118733_100000053191, *Caudoviricetes* IMGVR_UViG_3300049740_002374 histone ID: Ga0494453_0003361_6563_7693 and Ga0494453_0003361_7794_8051), the AlphaFold3 webserver (accessed on 2024/08) [38] was used. A range of candidate viral histones were used as input, along with the 147 bp W601 DNA sequence and its complementary reverse sequence (“ATCAGAATCCCGGTGCCGAGGCCGCTCAATTGGTCGTAGACAGCTCTAGCACCGCTTAAACGCACGTACGCGCTGTCCCCCGCGTTTTAACCGCCAAGGGGATTACTCCCTAGTCTCCAGGCACGTGTCAGATATATACATCGAT”) [25]. For viral genomes containing multiple histones, the corresponding histones were incorporated into the model according to their domain count, ensuring that a total of eight histone domains were represented in the model.

### Functional inference based on structural homology

For functional interference of viral histones and histone-fold proteins, we predicted their 3D structures using AlphaFold2, as mentioned above. For each protein structure, we searched for structural homologues against the AFDB [39] using FoldSeek [40] with default parameters, and annotated protein functions and/or functional domains. The structural matches e-value smaller than 0.001 were kept for further analysis. The matches were then ranked by the number of identical descriptions, and the top five non-redundant annotations were retained for each target protein.

### Histone similarity network and community detection

To cluster viral histones and histone-fold proteins, we extracted per-residue embeddings per sequence using the protein language model ProstT5 [41], and computed all-against-all embedding-based alignment (EBA) scores following the EBA protocol [42]. The protein sequence homology was determined by calculating all-against-all BLASTp v2.5.0+ [43] bit scores. Results were filtered based on the average EBA score (considering only those with maximum EBA scores of greater than 3.5) and the BLASTp p-value (considering only those with *p* less than 0.00001). The EBA scores and BLASTp average bit scores were normalized separately using min-max scaling. Then, the protein-protein similarities were determined using summed normalized EBA scores and BLASTp bit scores. The protein similarity network was built using Graphia version 5.1 [44] and clustered via Louvain weight clustering [45].

### Histone purification and nucleosome reconstitution

The coding DNA sequences of representative viral histone triplets and singlets were synthesized (GenScript Biotech) and cloned into pRSFDuet-1 vector. Each pair of viral histone triplet and singlet from the same viral genome were co-expressed in *E. coli* strain Rosetta (DE3). Cell cultures were grown to OD_600_ about 0.6 and induced with 0.4 mM isopropyl 1-thio-B-D-galactopyranoside for 4 h at 37 °C. Cells were pelleted and sonicated in buffer containing 20 mM Tris-HCl pH 8.0, 0.5 M NaCl, 0.1 mM EDTA, 1 mM DTT and PMSF. Supernatant was loaded onto a 5 mL HiTrap heparin column (Cytiva) and eluted with a gradient of 0.5 to 2 M NaCl. Fractions containing histone complexes were pooled and further purified by size exclusion chromatography in buffer containing 20 mM Tris-HCl pH 8.0, 2 M NaCl, 0.1 mM EDTA, 1 mM DTT. Samples were analyzed by SDS-PAGE and Coomassie-blue staining. Reconstitution of nucleosomes from purified viral histones were carried out by canonical salt gradient dialysis as described previously [25].

### Analysis of viral histone-fold proteins

For each protein, monomeric and different homo-oligomeric (dimeric, tetrameric, and octameric) structures were predicted using AlphaFold2 as mentioned above. The initial definition of histone-fold proteins clusters was conducted using HMM profile hits, the ProstT5 model and Louvain weighted clustering, as detailed above. Subsequently, structural comparisons of histone-fold proteins were conducted using ChimeraX [46], both within each cluster and between clusters, in order to calculate RMSD values. Furthermore, a range of known histones, including eukaryotic H2A, H2B, H3 and H4 (PDB ID 1KX5), archaeal histones (HMfA/B), bacterial histones (Bd0055), and ten recently identified types of prokaryotic histone-fold proteins [11], were used as references for structural comparisons to help annotate each cluster of viral histone-fold proteins, where appropriate oligomerization states were considered. When previously undescribed domain(s) were found in all members of a cluster or no known references could be properly used for annotations, the cluster was assigned a novel name. The results were further manually inspected using PyMOL v3.0.3 [The PyMOL Molecular Graphics System, Version 3.0 Schrödinger, LLC]. All figures related to 3D structures were generated using ChimeraX.

### Identification and analysis of chromatin-related proteins

We used a combination of previously reported chromatin-related domains from Grau-Bové X [47], and EpiFactors database version 2.1 [48]. The domains HMM profiles were downloaded from Pfam database [49]. HMMER was used to search for the viral proteins with chromatin-related domains. The chromatin-related proteins with e-values smaller than 0.001 were retained for further analysis.

For each IMG/VR viral Class, we examined correlation between histone presence (Hmmsearch e-value below 0.001) and number of genes that belong to each chromatin-related domain. We defined the presence (denoted as 1) of histones when there is at least one histone protein in the viral genome, and the absence (denoted as 0) as having none. For each viral class, we fitted a generalized linear model (GLM) with a binomial distribution and canonical link function, using histone presence or absence as the response variable, and the interaction between CRPs count and total protein number as predictors. To reduce sample size bias in the model, 1,000 histone-containing genomes and 1,000 histone-lacking genomes were randomly selected within each virus class, and GLM analysis was repeated 100 times. After 100 independent random samplings, we applied p-value adjustment using the Benjamini-Hochberg method and recorded both the minimum adjusted p-value and the number of significances (out of 100 random samplings) as a measurement of association strength. For *Caudoviricetes* and *Megaviricetes*, we applied the same model independently to each type of CRPs. The model performance was evaluated using AIC and McFadden’s pseudo-R square, and the statistical significance was determined using a *p*-value threshold of 0.05. The GLM were fit with glm() function from the *stats* package in R 4.3 (https://www.r-project.org/).

### Histone phylogenetic construction and viral host prediction

Phylogenetic trees of histone repeats were constructed based on their predicted tertiary structures using AlphaFold2. Briefly, these structures were aligned using Foldmason version 1.763a428 [50] easy-msa (default parameter: ––match-ratio 0.51 –– filter-msa 1 ––gap-open aa:10,nucl:10 ––gap-extend aa:1,nucl:1 ––precluster 1 ––report-mode 1). The alignment quality was manually inspected for 3Di alignment. The sequence alignment was trimmed using trimal (–gt 0.2 –resoverlap 0.5 –seqoverlap 50) [51], and the trimmed file was used for phylogenetic tree construction in IQ-TREE2 [52] with the model LG+C50+F+R7, employing 1000 ultrafast bootstrap replicates (–bb 1000) and 1000 SH-aLRT replicates (–alrt 1000), as previously described [18]. Viral hosts were predicted via iPHoP using iPHoP_db_Aug23_rw database [53]. The host tree was curated using GTDB tree release220 [54].

## Data availability

The metadata of predicted histones, histone-fold proteins and cluster IDs have been summarized in the supplementary tables. Supplementary tables, viral histone(–fold) similarity networks, phylogenetic trees, predicted viral histones(–fold) structures, predicted viral nucleosome core particle structures, and codes of GLM analysis are available on Figshare under the DOI: 10.6084/m9.figshare.27950739.

## Acknowledgements

This work was funded by the National Key Research and Development Program of China (2023YFC2306800 to Yue Liu) and the National Natural Science Foundation of China (82188102 to Haibo Wang).

## Author contribution

Yang Liu: writing – review & editing, writing – original draft, bioinformatics analysis, conceptualization. Zhuru Hou: writing – review & editing, writing – original draft, bioinformatics analysis. Wanshan Hao: writing – review & editing, biological experiments. Shaoqing Cui: writing – review & editing. Haibo Wang: writing – review & editing, supervision, funding acquisition, conceptualization. Yue Liu: writing – review & editing, writing – original draft, supervision, funding acquisition, conceptualization.

