## Supplementary_figures for "Metagenomic mining reveals novel viral histones in dsDNA viruses"

### Supplementary Figure Legends

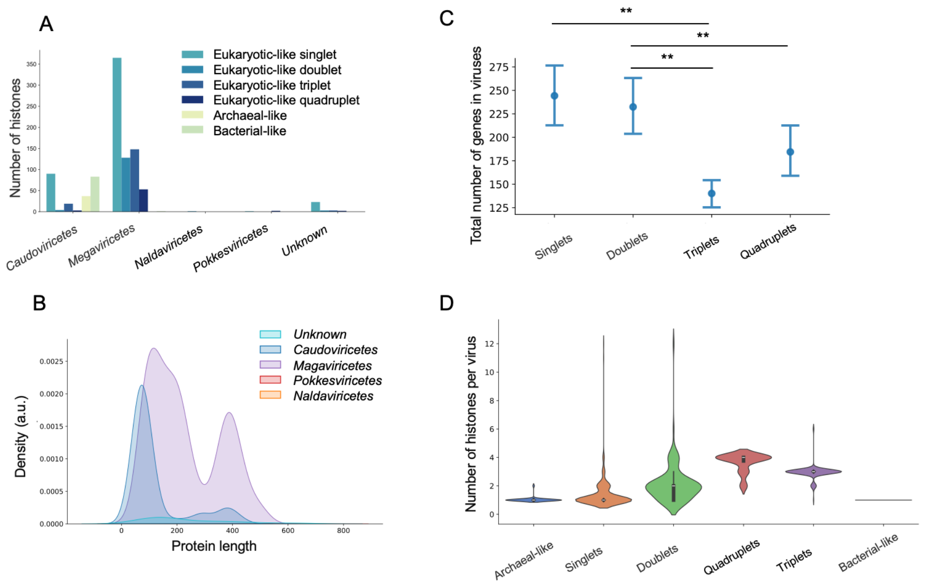


**Figure S1. Viral histones from the IMG/VR database.** (A) Abundance of different viral histone types across two viral classes, where histone types are annotated and colored by their corresponding HMM profiles. (B) Histone protein length distribution across virus classes. (C) The total number of genes in eukaryotic-like histone-containing viruses is represented with confidence intervals. Statistical significance was determined using the Mann-Whitney U test with $p < 0.05$. (D) The total number of histones identified in every viral genome categorized and colored based on different histone types.

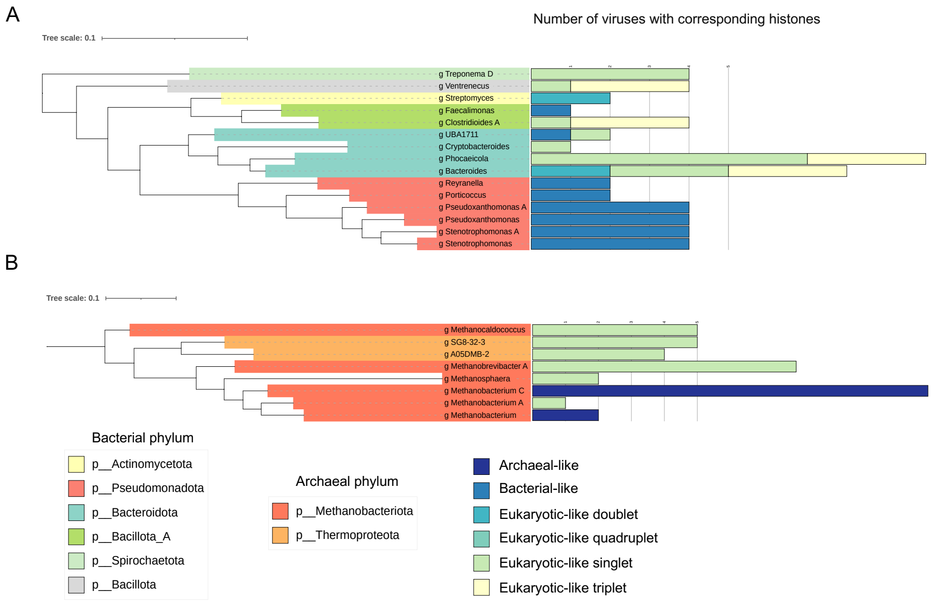


**Figure S2. Host predictions for histone-containing viral genomes within *Caudoviricetes*.** (A) The bacterial phylogenetic tree of predicted viral hosts for the *Caudoviricetes* class. The abundance of viruses color-coded based on the types of histones they contain. Phylogenetic topology and distances were obtained from the GTDB database (release 220). (B) The archaeal phylogenetic tree of predicted viral hosts for *Caudoviricetes* class.

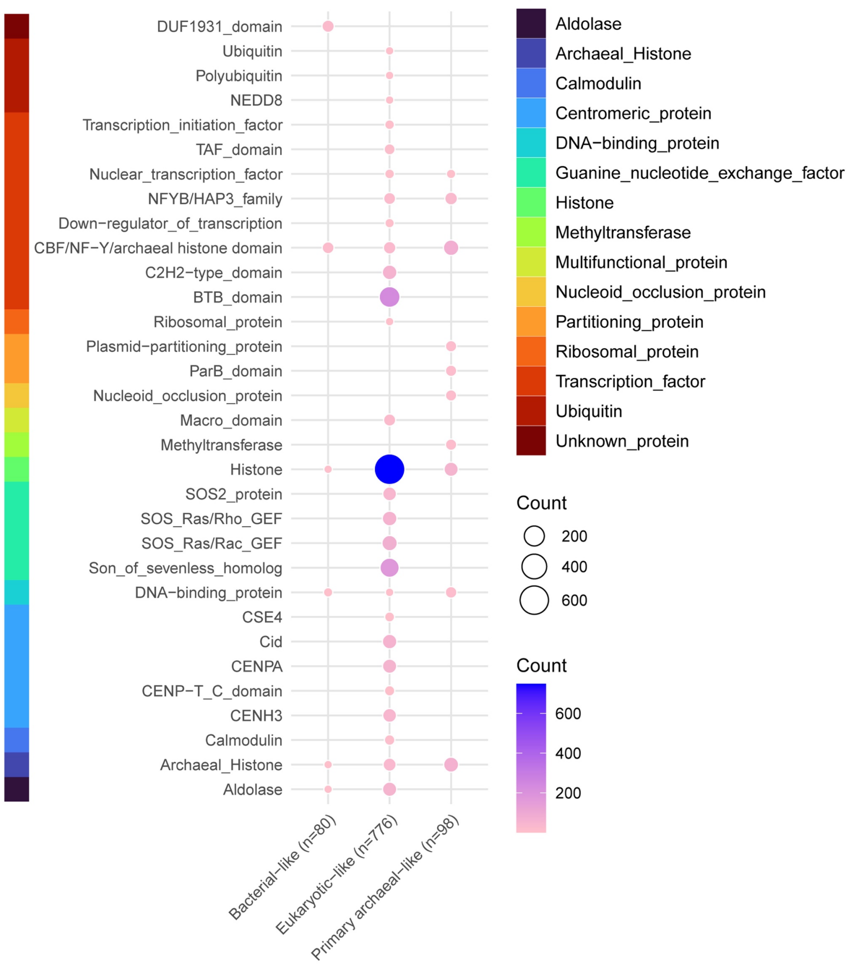


**Figure S3. Structure-based functional domain annotations of identified viral histones.** The x-axis represents the different classifications of viral histones, while the y-axis indicates their predicted functional domains. The circles represent the degree of enrichment of these histones.

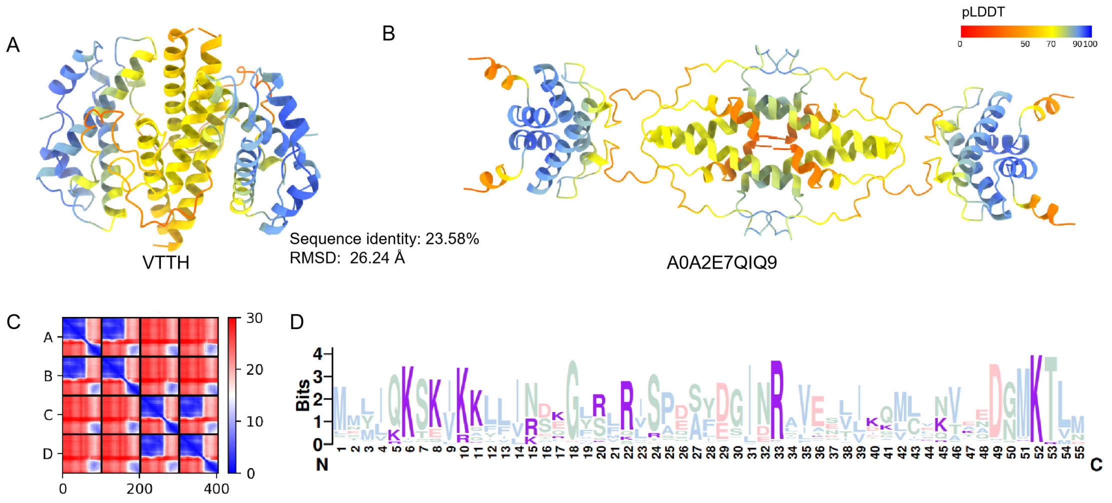

**Figure S4. The structural characteristics of VTTHs.** (A-B) The tetrameric structures of a typical VTTH (A) and of phage histone A0A2E7QIQ9 used as a reference for HMM-based identification of VTTHs (B), as predicted by AlphaFold2. (C) Predicted aligned error plots for the VTTH tetramer predictions. (D) Logo representation of all identified VTTHs.

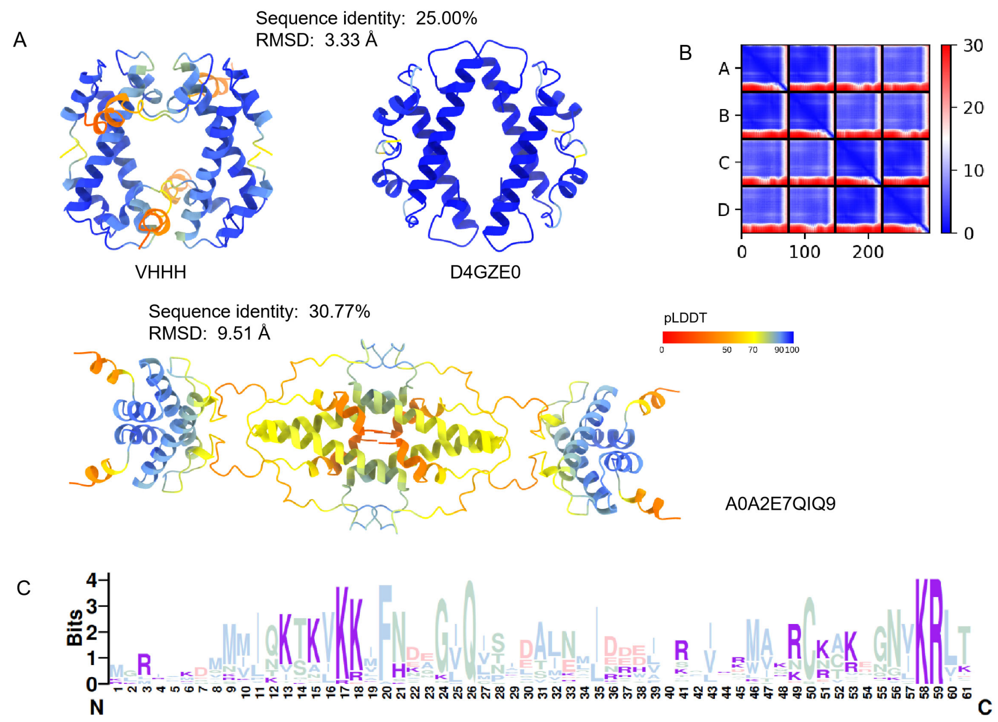


**Figure S5. The structural characteristics of VHHHs.** (A) The tetrameric structures of a representative VHHH, and prokaryotic histone A0A2E7QIQ9 (used as a reference for HMM-based identification of VHHHs) and face-to-face prokaryotic histone D4GZE0 (with which VHHHs share structural similarities), as predicted by AlphaFold2. (B) Predicted aligned error plots for the VHHH tetramer predictions. (C) Logo representation of all identified VHHHs.

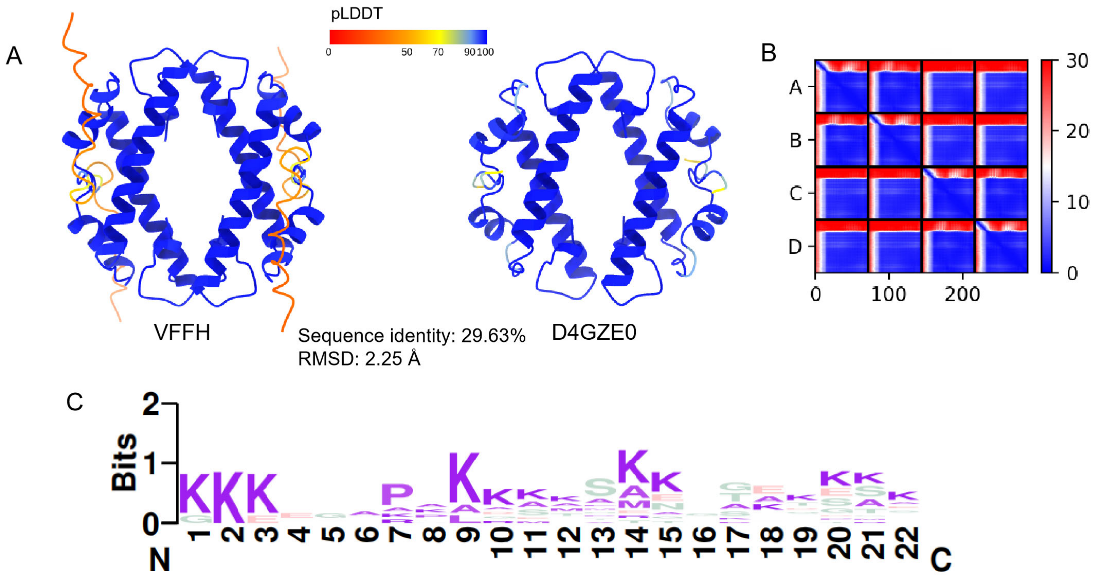


**Figure S6. The structural characteristics of VFFHs.** (A) The predicted tetrameric structure of a typical VFFH and face-to-face prokaryotic histone D4GZE0 (used as a reference for HMM-based identification of VFFHs). (B) Predicted aligned error plots for the VFFH tetramer predictions. (C) Logo representation of N-terminal amino acid sequences for VFFHs.

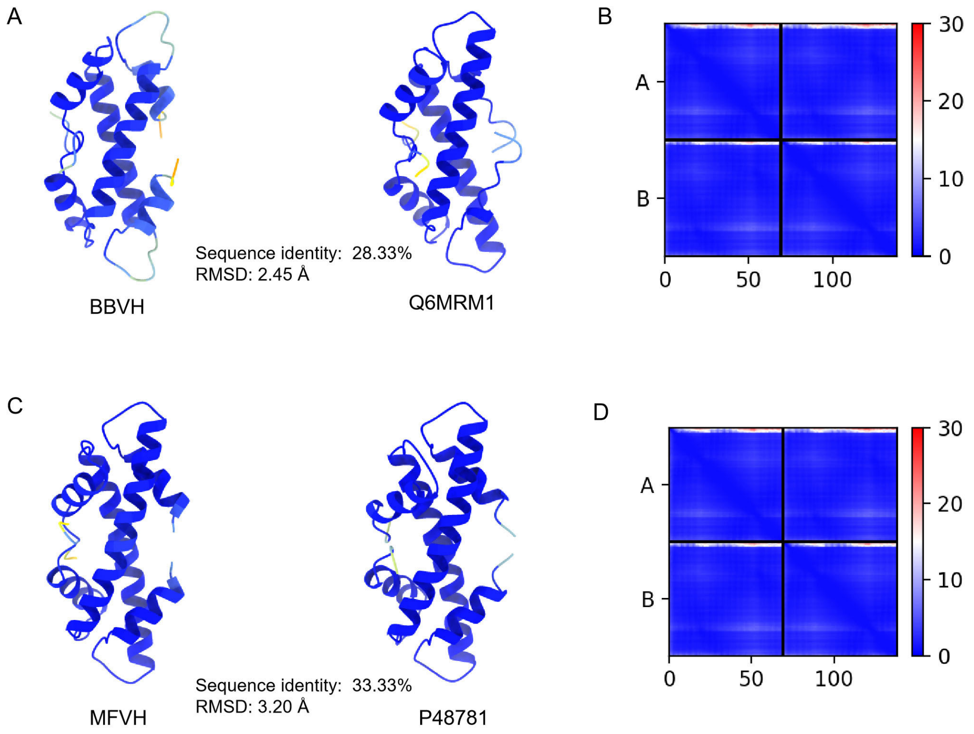

**Figure S7. The structural characteristics of BBVHs and MFVHs.** (A) The dimeric structures of a typical BBVH and *Bdellovibrio bacteriovorus* Bd0055 (Q6MRM1, with which BBVHs share structural similarities) as predicted by AlphaFold2. (B) Predicted aligned error plots for the BBVH dimer predictions. (C) The dimeric structures of a typical MFVH and *Methanothermus fervidus* HmfA (P48781, with which MFVHs share structural similarities) as predicted by AlphaFold2. (D) Predicted aligned error plots for the MFVH dimer predictions.

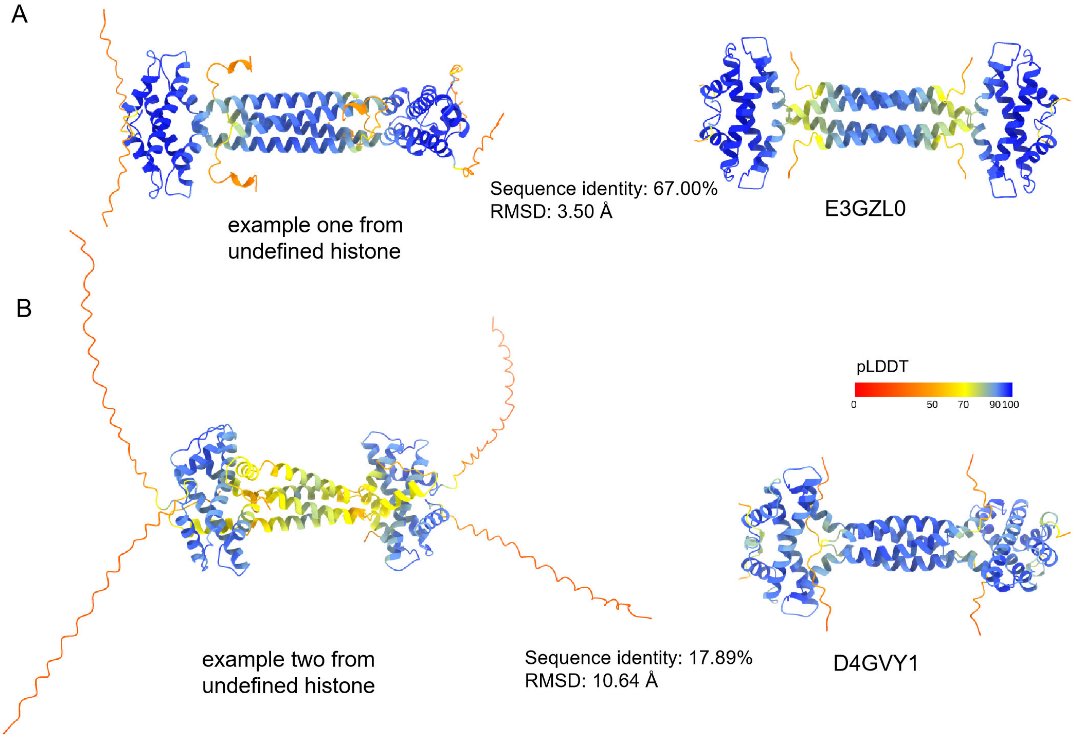


**Figure S8. The structural characteristics of undefined viral histone-fold proteins.** Shown are two examples in comparison with the prokaryotic histones coiled-coil histone E3GZL0 (A) and RdgC histone D4GVY1 (B) structures, respectively.

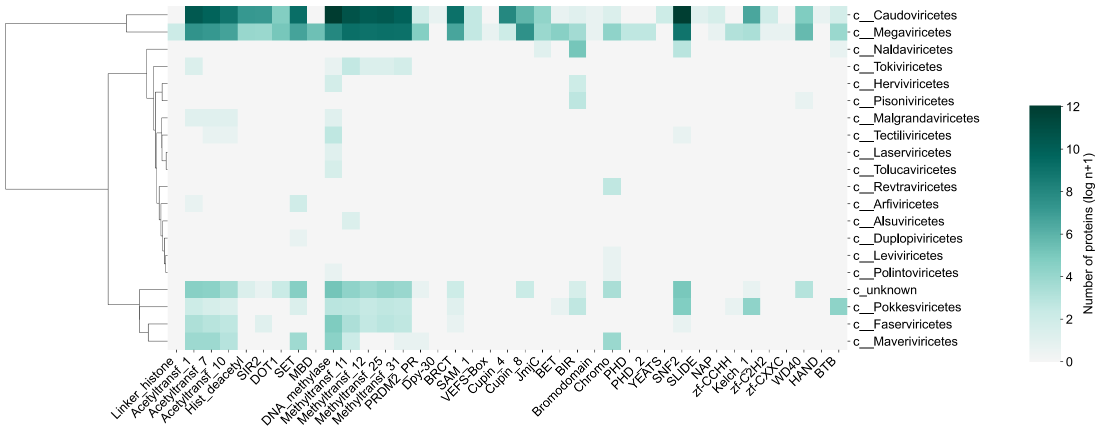


**Figure S9. The prevalence of chromatin-related proteins identified from different virus classes.** The viral class was hierarchically clustered using the Ward D2 method based on abundance pattern.

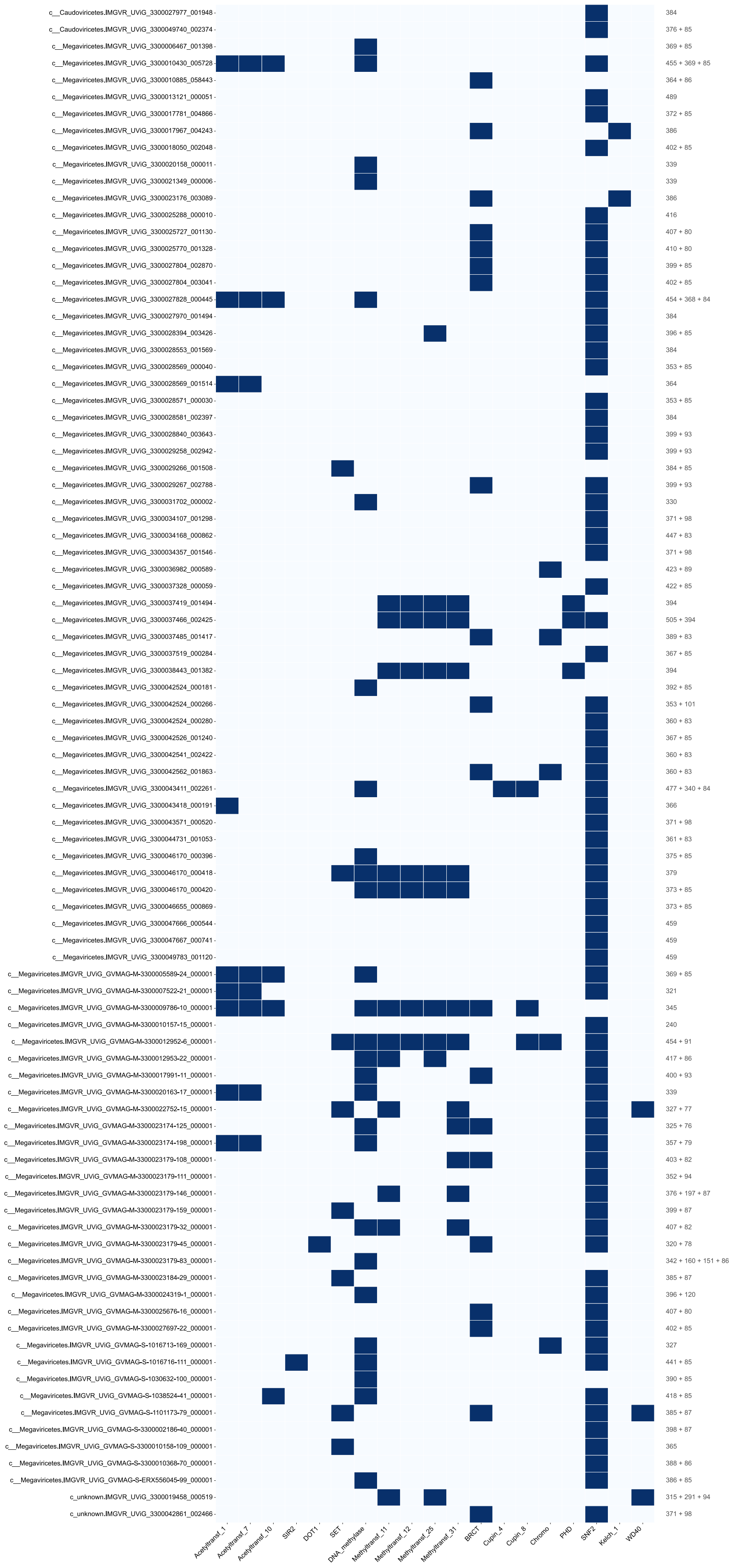

c__Caudoviricetes.IMGVR_UViG_3300027977_001948
c__Caudoviricetes.IMGVR_UViG_3300049740_002374
c__Megaviricetes.IMGVR_UViG_3300006467_001398
c__Megaviricetes.IMGVR_UViG_3300010430_005728
c__Megaviricetes.IMGVR_UViG_3300010885_058443
c__Megaviricetes.IMGVR_UViG_3300013121_000051
c__Megaviricetes.IMGVR_UViG_3300017781_004866
c__Megaviricetes.IMGVR_UViG_3300017967_004243
c__Megaviricetes.IMGVR_UViG_3300018050_002048
c__Megaviricetes.IMGVR_UViG_3300020158_000011
c__Megaviricetes.IMGVR_UViG_3300021349_000006
c__Megaviricetes.IMGVR_UViG_3300023176_003089
c__Megaviricetes.IMGVR_UViG_3300025288_000010
c__Megaviricetes.IMGVR_UViG_3300025727_001130
c__Megaviricetes.IMGVR_UViG_3300025770_001328
c__Megaviricetes.IMGVR_UViG_3300027804_002870
c__Megaviricetes.IMGVR_UViG_3300027804_003041
c__Megaviricetes.IMGVR_UViG_3300027828_000445
c__Megaviricetes.IMGVR_UViG_3300027970_001494
c__Megaviricetes.IMGVR_UViG_3300028394_003426
c__Megaviricetes.IMGVR_UViG_3300028553_001569
c__Megaviricetes.IMGVR_UViG_3300028569_000040
c__Megaviricetes.IMGVR_UViG_3300028569_001514
c__Megaviricetes.IMGVR_UViG_3300028571_000030
c__Megaviricetes.IMGVR_UViG_3300028581_002397
c__Megaviricetes.IMGVR_UViG_3300028840_003643
c__Megaviricetes.IMGVR_UViG_3300029258_002942
c__Megaviricetes.IMGVR_UViG_3300029266_001508
c__Megaviricetes.IMGVR_UViG_3300029267_002788
c__Megaviricetes.IMGVR_UViG_3300031702_000002
c__Megaviricetes.IMGVR_UViG_3300034107_001298
c__Megaviricetes.IMGVR_UViG_3300034168_000862
c__Megaviricetes.IMGVR_UViG_3300034357_001546
c__Megaviricetes.IMGVR_UViG_3300036982_000589
c__Megaviricetes.IMGVR_UViG_3300037328_000059
c__Megaviricetes.IMGVR_UViG_3300037419_001494
c__Megaviricetes.IMGVR_UViG_3300037466_002425
c__Megaviricetes.IMGVR_UViG_3300037485_001417
c__Megaviricetes.IMGVR_UViG_3300037519_000284
c__Megaviricetes.IMGVR_UViG_3300038443_001382
c__Megaviricetes.IMGVR_UViG_3300042524_000181
c__Megaviricetes.IMGVR_UViG_3300042524_000266
c__Megaviricetes.IMGVR_UViG_3300042524_000280
c__Megaviricetes.IMGVR_UViG_3300042526_001240
c__Megaviricetes.IMGVR_UViG_3300042541_002422
c__Megaviricetes.IMGVR_UViG_3300042562_001863
c__Megaviricetes.IMGVR_UViG_3300043411_002261
c__Megaviricetes.IMGVR_UViG_3300043418_000191
c__Megaviricetes.IMGVR_UViG_3300043571_000520
c__Megaviricetes.IMGVR_UViG_3300044731_001053
c__Megaviricetes.IMGVR_UViG_3300046170_000396
c__Megaviricetes.IMGVR_UViG_3300046170_000418
c__Megaviricetes.IMGVR_UViG_3300046170_000420
c__Megaviricetes.IMGVR_UViG_3300046655_000869
c__Megaviricetes.IMGVR_UViG_3300047666_000544
c__Megaviricetes.IMGVR_UViG_3300047667_000741
c__Megaviricetes.IMGVR_UViG_3300049783_001120
c__Megaviricetes.IMGVR_UViG_GVMAG-M-3300005589-24_000001
c__Megaviricetes.IMGVR_UViG_GVMAG-M-3300007522-21_000001
c__Megaviricetes.IMGVR_UViG_GVMAG-M-3300009786-10_000001
c__Megaviricetes.IMGVR_UViG_GVMAG-M-3300010157-15_000001
c__Megaviricetes.IMGVR_UViG_GVMAG-M-3300012952-6_000001
c__Megaviricetes.IMGVR_UViG_GVMAG-M-3300012953-22_000001
c__Megaviricetes.IMGVR_UViG_GVMAG-M-3300017991-11_000001
c__Megaviricetes.IMGVR_UViG_GVMAG-M-3300020163-17_000001
c__Megaviricetes.IMGVR_UViG_GVMAG-M-3300022752-15_000001
c__Megaviricetes.IMGVR_UViG_GVMAG-M-3300023174-125_000001
c__Megaviricetes.IMGVR_UViG_GVMAG-M-3300023174-198_000001
c__Megaviricetes.IMGVR_UViG_GVMAG-M-3300023179-108_000001
c__Megaviricetes.IMGVR_UViG_GVMAG-M-3300023179-111_000001
c__Megaviricetes.IMGVR_UViG_GVMAG-M-3300023179-146_000001
c__Megaviricetes.IMGVR_UViG_GVMAG-M-3300023179-159_000001
c__Megaviricetes.IMGVR_UViG_GVMAG-M-3300023179-32_000001
c__Megaviricetes.IMGVR_UViG_GVMAG-M-3300023179-45_000001
c__Megaviricetes.IMGVR_UViG_GVMAG-M-3300023179-83_000001
c__Megaviricetes.IMGVR_UViG_GVMAG-M-3300023184-29_000001
c__Megaviricetes.IMGVR_UViG_GVMAG-M-3300024319-1_000001
c__Megaviricetes.IMGVR_UViG_GVMAG-M-3300025676-16_000001
c__Megaviricetes.IMGVR_UViG_GVMAG-M-3300027697-22_000001
c__Megaviricetes.IMGVR_UViG_GVMAG-S-1016713-169_000001
c__Megaviricetes.IMGVR_UViG_GVMAG-S-1016716-111_000001
c__Megaviricetes.IMGVR_UViG_GVMAG-S-1030632-100_000001
c__Megaviricetes.IMGVR_UViG_GVMAG-S-1038524-41_000001
c__Megaviricetes.IMGVR_UViG_GVMAG-S-1101173-79_000001
c__Megaviricetes.IMGVR_UViG_GVMAG-S-3300002186-40_000001
c__Megaviricetes.IMGVR_UViG_GVMAG-S-3300010158-109_000001
c__Megaviricetes.IMGVR_UViG_GVMAG-S-3300010368-70_000001
c__Megaviricetes.IMGVR_UViG_GVMAG-S-ERX556045-99_000001
c_unknown.IMGVR_UViG_3300019458_000519
c_unknown.IMGVR_UViG_3300042861_002466
384
376 + 85
369 + 85
455 + 369 + 85
364 + 86
489
372 + 85
386
402 + 85
339
339
386
416
407 + 80
410 + 80
399 + 85
402 + 85
454 + 368 + 84
384
396 + 85
384
353 + 85
364
353 + 85
384
399 + 93
399 + 93
384 + 85
399 + 93
330
371 + 98
447 + 83
371 + 98
423 + 89
422 + 85
394
505 + 394
389 + 83
367 + 85
394
392 + 85
353 + 101
360 + 83
367 + 85
360 + 83
360 + 83
477 + 340 + 84
366
371 + 98
361 + 83
375 + 85
379
373 + 85
373 + 85
459
459
459
369 + 85
321
345
240
454 + 91
417 + 86
400 + 93
339
327 + 77
325 + 76
357 + 79
403 + 82
352 + 94
376 + 197 + 87
399 + 87
407 + 82
320 + 78
342 + 160 + 151 + 86
385 + 87
396 + 120
407 + 80
402 + 85
327
441 + 85
390 + 85
418 + 85
385 + 87
398 + 87
365
388 + 86
386 + 85
315 + 291 + 94
371 + 98
Acetyltransf_1
Acetyltransf_7
Acetyltransf_10
SIR2
DOT1
SET
DNA_methylase
Methyltransf_11
Methyltransf_12
Methyltransf_25
Methyltransf_31
BRCT
Cupin_4
Cupin_8
Chromo
PHD
SNF2
Kelch_1
WD40

**Figure S10. The presence or absence chromatin-related proteins on histone triplet-containing viral genomes.** The text on the right of each row indicates the length combinations of multiple co-occurring histones in the viral genomes. Only viral histone triplet-containing genomes with at least one putative chromatin-related protein are included in this figure.

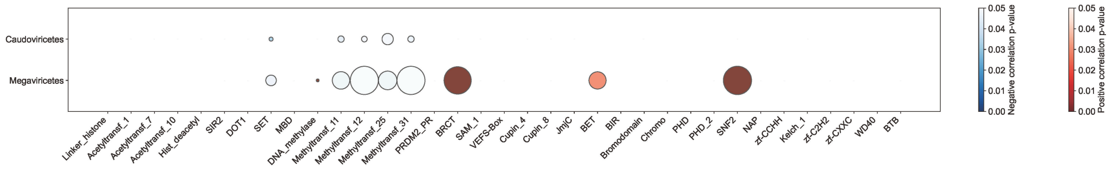


**Figure S11. The statistical correlation between the number of putative** chromatin-related proteins and the presence of histones on the viral genomes. The positive significant correlation is indicated by red ($p < 0.05$), and negative significant correlation is indicated by blue ($p < 0.05$). The sizes of the circle indicate the number of significant observations out of 100 independent random samplings. Blank indicates a lack of significant correlation.

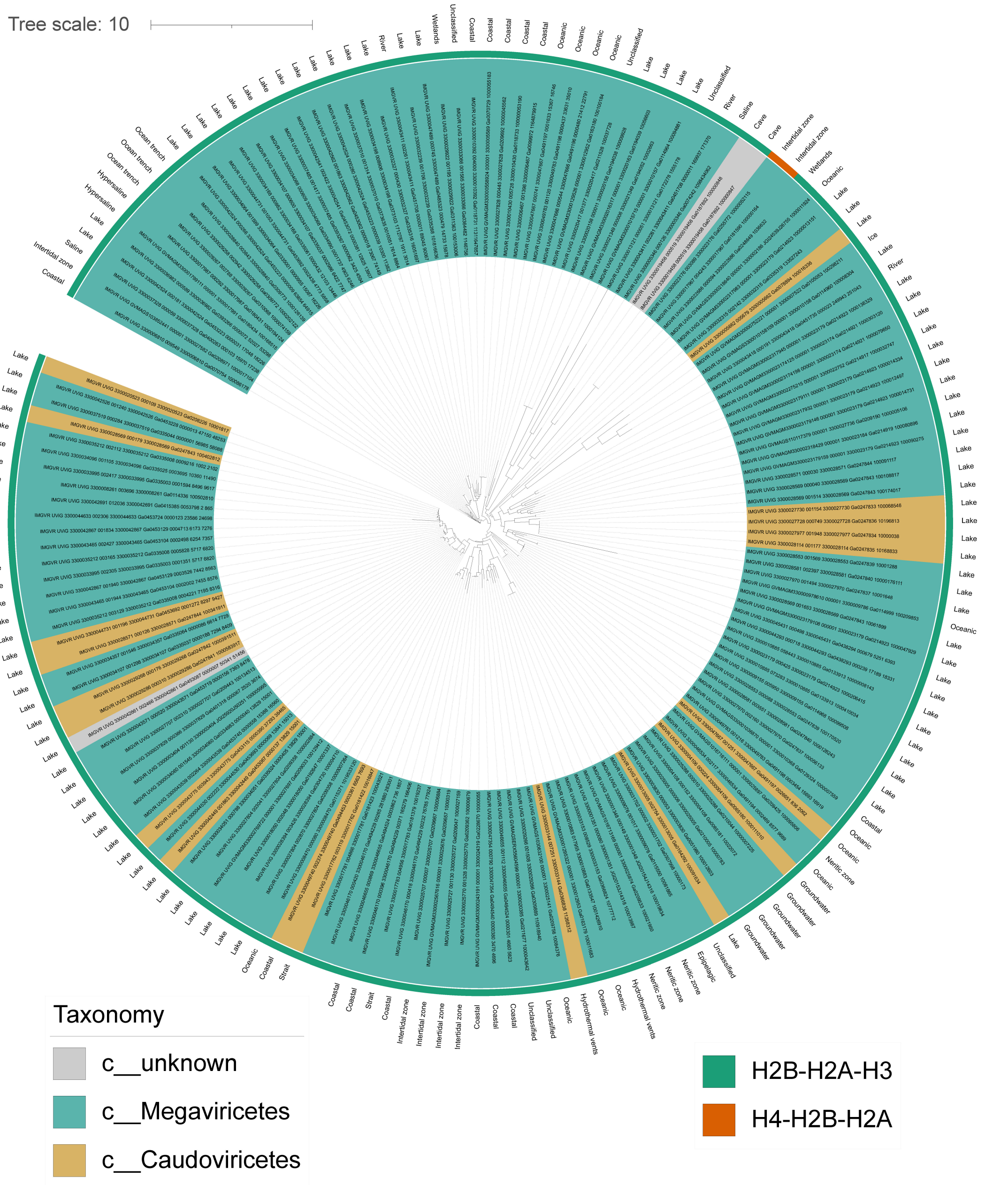


**Figure S12. The histone triplet structure-based phylogenetic tree.** Structure-based phylogenetic reconstruction of identified histone triplets (see Method). The taxonomy and habitats of correspond viruses are labeled according to IMG/VR metadata.
